# Using deep amplicon sequencing as a molecular xenomonitoring approach for detecting filarial nematodes in biting arthropod vectors

**DOI:** 10.1101/2025.07.29.667378

**Authors:** Matthew R. Kulpa, Chris Blazier, John S. Gileard, Guilherme G. Verocai

**Affiliations:** Department of Veterinary Pathobiology, College of Veterinary Medicine and Biomedical Sciences, Texas A&M University, College Station, Texas, United States of America; Texas A&M Institute of Genome Sciences and Society, Texas A&M University, College Station, Texas, United States of America; Department of Comparative Biology and Experimental Medicine, Faculty of Veterinary Medicine, University of Calgary, Calgary, Alberta, Canada

**Keywords:** Filarioidea, Lymphatic filariasis, Molecular xenomonitoring, Neglected Tropical Diseases, One Health, Oxford Nanopore, Tropical medicine, Vector-borne pathogens

## Abstract

Filarial nematodes are an important group of parasites that impact public, veterinary, and wildlife health globally. In order to understand these impacts and minimize their effects, scientists use molecular xenomonitoring techniques to understand their distribution and track elimination efforts. However, these molecular techniques can have narrow diagnostic capacity due to their species-specific approach which limits our understanding of important co-endemic filarial nematodes. Next-generation sequencing offers the ability to detect multiple species of coinfecting filarial nematodes and thus improve are ability to monitor, treat, and eliminate these pathogens. In this paper, we have developed a deep amplicon sequencing approach using filarial nematode primers targeting the cytochrome oxidase c subunit 1 (*coxI*) gene. To replicate molecular xenomonitoring conditions, third stage larvae (L3) of three species of filarioid nematodes (*Brugia malayi*, *Brugia pahangi*, *Dirofilaria immitis*) were spiked in different proportions to pools comprising various amounts of female *Aedes aegypti* mosquitoes (0, 10, 50, 100). Each pool was subjected to DNA extraction and Oxford Nanopore Technologies (ONT) deep amplicon sequencing protocols. Two sets of demultiplexing pipelines were utilized to optimize this novel approach, each reaching, 92.71% and 97.92% accuracy in identification of species composition across mock pools. However, in heterogenous pools, filarial species *D. immitis* exhibited an overrepresentation of reads and *B. pahangi* an underrepresentation of reads. We discuss reasons for recount biases and how this new molecular xenomonitoring tool could be implemented to serve public health, veterinary medicine, and scientific advancement.

Note: Supplementary data associated with this article

**Author summary:** Human and animal diseases caused by filarial nematodes affect millions of people worldwide, particularly in low-income countries. These parasites are transmitted by blood-feeding arthropod vectors, such as mosquitoes and black flies. Thus, a major sector of public health research focuses on how to monitor, treat, and eliminate these harmful pathogens. An effective way is to capture these arthropod vectors and molecularly test these for filarial DNA in large sample pools. However, these pools can comprise multiple filarial species which targeted genetic analysis can miss. This proof-of-concept study seeks to circumvent these issues by using new next-generation sequencing approaches to capture the wider filarial diversity that may be contained in a single vector pool. We believe this tool could largely be beneficial to governments and organizations seeking to eliminate these filarial nematodes and become certified as a region free of certain devastating filarial species. Furthermore, we know very little about filarial diversity and this could be an integral tool to define their geographic distribution and future emerging threats to both human and animal health.

## Introduction

Filarial nematodes are vector-borne parasites that are of global significance to human health. Many filarial nematodes are agents for neglected tropical diseases (NTDs) in humans such as river blindness (e.g., *Onchocerca volvulus* (Leuckart, 1893)) and lymphatic filariases (e.g., *Wuchereria bancrofti* (Cobbold, 1877), *Brugia malayi* (Brug, 1927) and *Brugia timori* Partono, 1977) [1]. The World Health Organization (WHO) currently estimates that 25 million people are infected by *O. volvulus*, with another 90 million at risk and 51 million people are infected with parasitic agents causing lymphatic filariases with another 657 million at risk [2, 3]. Molecular xenomonitoring, by collecting hematophagous arthropods to screen for pathogen DNA, is a primary tool for detecting these important filarial worms in endemic areas [4]. WHO guidelines for combating filarial infections in endemic regions are largely based on mass drug administration efforts, which rely on molecular xenomonitoring surveillance for tracking parasite’s epidemiology and certifying local or in-country elimination [5–10]. The implementation of molecular surveillance has many advantages when compared to former blunt arthropod dissection techniques including: its higher-throughput capability, increased sensitivity, and ability to diagnose larval stages of filarial nematodes at the species-level. However, currently used molecular xenomonitoring tools such as conventional PCR or real-time PCR are often targeted and species-specific, which can be problematic in areas of co-endemicity for multiple filarial nematode s. For example, *O*. *volvulus* and *W*. *bancrofti* are two, among many, filarial nematodes found in Africa that infect humans and cause devastating disease [11]. Thus, targeted molecular approaches may overlook valuable public health information, which can lead to poor and costly decision-making regarding treatment and elimination efforts. For example, some African countries are co-endemic to both *O*. *volvulus* and *Loa loa* (Cobbold, 1864), the African eyeworm, with co-infections being possible. Implementation of mass drug administration with ivermectin targeting onchocerciasis can cause severe adverse reaction in co-infected individuals with high *L*. *loa* microfilaremia [12, 13].

Filarial nematode surveillance via arthropod vectors can yield not only data relevant to human health but may also provide insights into filarial species that infect domestic and wild animals. These can be in concern to companion animal veterinary medicine (e.g., *Cercopithifilaria bainae* Almeida & Vicente, 1984, *Dirofilaria immitis* (Leidy, 1856), *Onchocerca lupi* Rodonaja, 1967) wildlife health or conservation (e.g., *Elaeophora schneideri* Wehr & Dikmans, 1935, *Rumenfilaria andersoni* Lankester & Snider, 1982, and *Setaria tundra* Desset, 1966), and livestock health and production (e.g., *Onchocerca gibsoni* Cleland & Johnston, 1910, *Onchocerca ochengi* Bwangomoi, 1969, and *Parafilaria bovicola* Tubangui, 1934) [1, 14–16]. While most of these filarial nematodes are mainly associated with non-human hosts, many are zoonotic (e.g., *D*. *immitis*, *O*. *lupi)* [17–19]. Therefore, enhanced monitoring of domestic and wild animal filarial nematodes can be mutually beneficial to public health. In addition, recent evidence has shown extensive cryptic diversity in several genera of filarial nematodes and small genetic discrepancies can prevent proper diagnosis when using a targeted molecular diagnostic approach [20–23]. Exploration of these diverse populations or species can shed light on unknown host and vector associations, geographic distribution, and, possibly, mutations conferring drug resistance. Overall, the current model of molecular xenomonitoring for filarial nematodes does not fully capture the available information in a sample.

If a novel molecular diagnostic approach were developed to survey filarial nematodes co-occurring within vectors or other biological samples from vertebrate hosts, we would enhance molecular xenomonitoring capabilities and optimize resources, while concurrently broadening the One Health applications of surveillance. Continuous advancements in molecular analysis, such as next-generation sequencing, offer expanded parasite diagnostic capabilities. In fact, deep amplicon sequencing has been successfully employed to analyze the community of gastrointestinal nematodes in various domestic and wild mammalian hosts [24–31]. The approach, coined the “nemabiome” is analogous to how scientists explore microbial communities (i.e., microbiome), but most commonly utilizes the nematode ITS-2 rDNA locus instead of the 16S rDNA. We propose to extrapolate these high-throughput, deep amplicon sequencing methodologies to target DNA of filarial nematodes in arthropod vectors. A recent published work [32] describes amplification and deep sequencing of the cytochrome c oxidase subunit 1 (*cox1*) gene can identify different species of filarial nematodes. This molecular approach, the “filariome”, would be increasingly relevant for monitoring co-endemic filarial nematodes in pooled arthropod vectors throughout disease elimination efforts.

Improved filarial nematode monitoring through deep amplicon sequencing will rely on validation and optimization of these molecular techniques. Therefore, we explored the approach by creating ‘mock pools’ comprising mosquito heads and different species of infective third-stage larvae (L3s) from which to prepare template for deep amplicon sequencing to assess the accuracy of the method in determining the relative abundance of different filarial species during molecular xenomonitoring. We discuss the potential use of this comprehensive approach into surveillance of neglected tropical diseases, as relevant epidemiological data on related filarial nematodes associated with domestic animals and wildlife, may be of zoonotic, veterinary and conservation relevance may be effectively and simultaneously acquired.

## Methods

### Mock pool creation

Mock pools were created to replicate molecular xenomonitoring conditions. These pools were formed by using different numbers of heads from female *Aedes aegypti* mosquitoes, with various combinations of filarial nematode third-stage larvae (L3) spiked into each pool (Fig. 1, 2). Only competent arthropod vectors, or ones that can actively transmit filarial nematodes to a new vertebrate host, will contain L3 larvae within the head or mouth parts [4, 33]. Therefore, using only the head during molecular xenomonitoring, eliminates the possibility of accidental ingestion of filarial DNA by an incompetent arthropod vector. Three species of filarial nematodes of relevance to human and veterinary medicine were used: *B. malayi* (Bm), *Brugia pahangi* (Buckley & Edeson, 1956) (Bp), and *D. immitis* (Di). Mosquitoes were acquired from the Filariasis Research Reagent Resource Center (FR3) [34].

**Figure 1.**
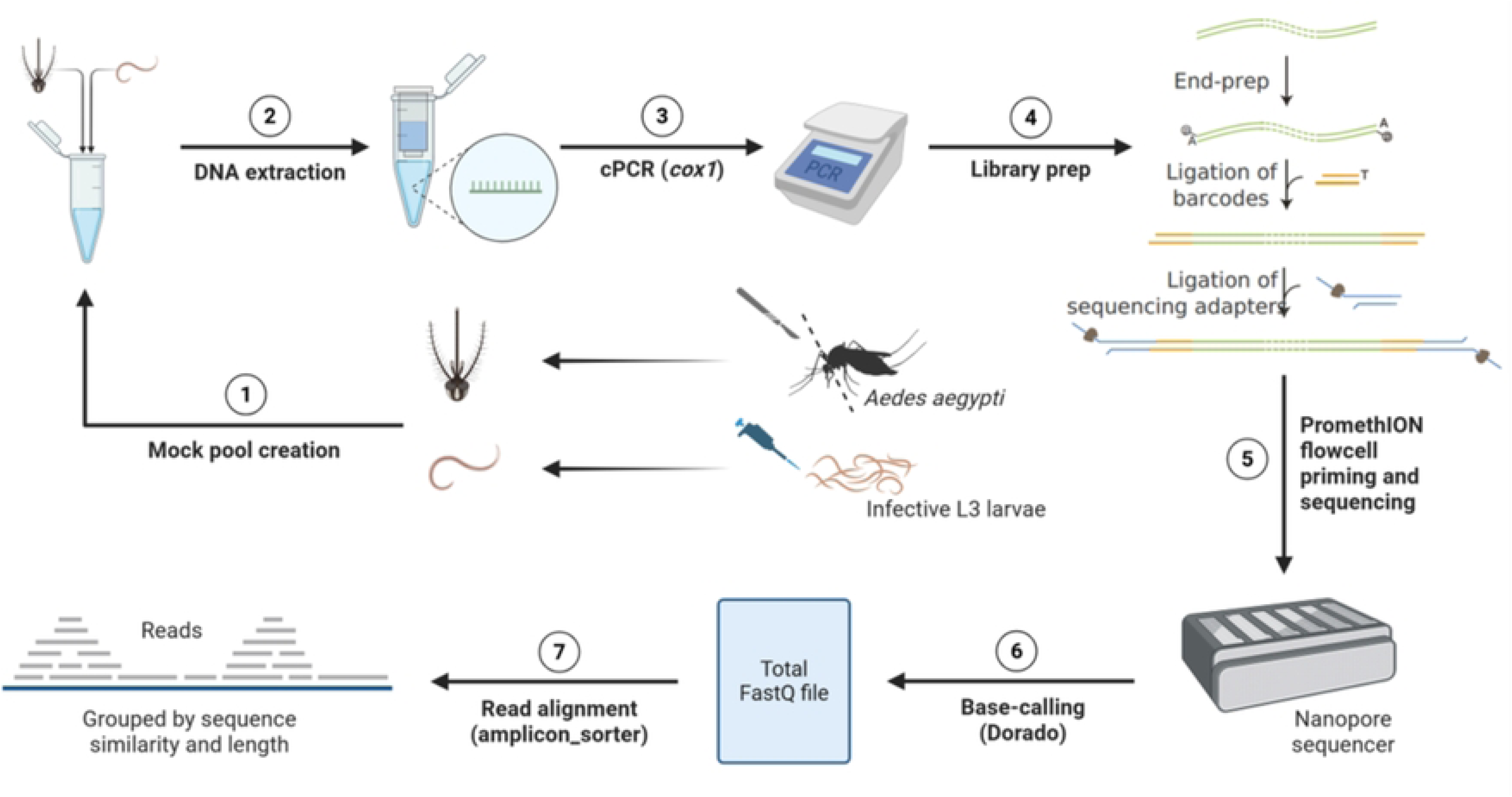
Workflow for using deep amplicon sequencing as a pan-filarial detection method for molecular xenomonitoring (i.e., filariome). (1) Mock pools were created by combining female *Aedes aegypti* mosquito heads with filarial nematodes (i.e., *Brugia malayi*, *Brugia pahangi*, and *Dirofilaria immitis*) in a bead homogenizer tube comprising 180 µL of TLA lysis buffer. (2) Physical break down of mosquito heads and worms were done using an Omni Bead Ruptor and enzymatically disrupted using Proteinase K (20 µL) and Thermomixer. Lysate was then added to Maxwell^®^ RSC Instrument and Tissue DNA Kit, where DNA was automatically processed for DNA extraction. (3) Conventional PCR was used to amplify *cox1* regions of filarial DNA. (5) Amplified material was then library prepped via a Native Barcoding Kit 96 V14, which added unique barcodes to each sample and ligated adapters. (6) Library prepped samples were then loaded into PromethION Nanopore sequencer where it was primed and sequenced. (7) Base-calling was performed using Dorado (8) and trimming and read alignment was done using Porechop and amplicon_sorter.

Mock pools were created using either 0, 10, 50, or 100 mosquito heads. These quantities mirror current molecular xenomonitoring surveillance efforts for monitoring and eliminating neglected tropical diseases, including lymphatic filariasis [4]. The “zero” mosquito pools allowed for both positive and negative controls. Mosquito heads were bluntly dissected from bodies using probes and a dissection microscope and added to their corresponding pools (Fig. 1). Once dissected, L3 belonging to each species of filarial nematode were then spiked (or not spiked) into each mosquito head pool in quantities of 0, 1, 5, and 10 L3 via a 1 µL pipette (Fig. 2). Overall, 64 variations of filarial species, and ratio of L3 and mosquito heads were established and done in triplicate to produce a 192 mock pool samples (Table 1).

**Figure 2.**
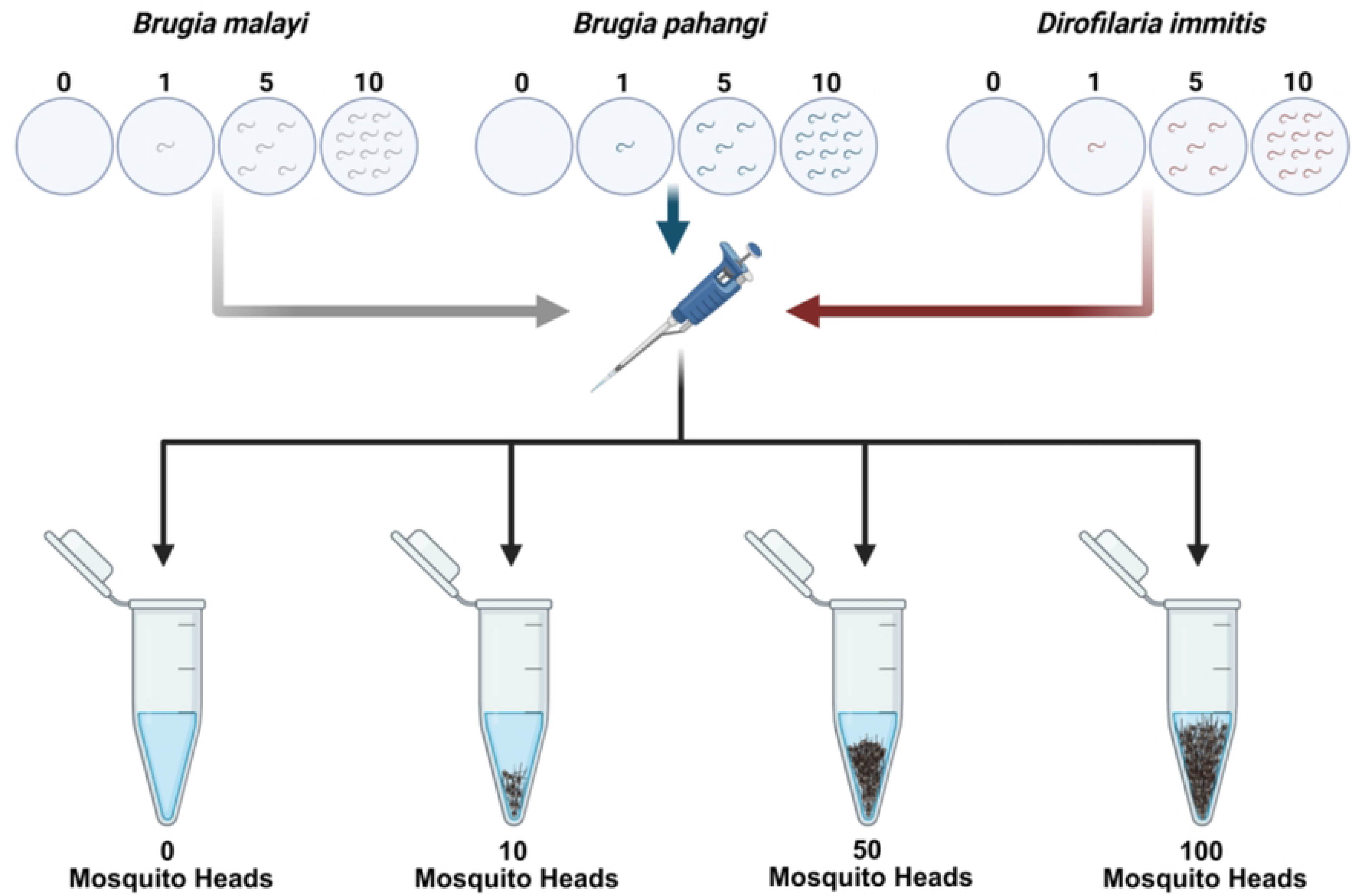
Mock pool design and composition. Mock pools were created by first removing the heads of female *Aedes aegypti* mosquitoes from their body. These heads were then added (or not added) to a bead homogenizer tube in quantities of 0, 10, 50, 100 depending on the sample. Each bead homogenizer tube comprised 180 µL of TLA lysis buffer. After dissection, three different species of infective L3 larvae (i.e., *Brugia malayi*, *Brugia pahangi*, and *Dirofilaria immitis*) were individually pipetted into these bead homogenizer tubes in quantities of 0, 1, 5, or 10. Overall, 64 variations of filarial species and/or mosquito heads were created in triplicate for a total of a 192 mock pools. Specific tube details and their contents can be found in Table 1. Each mock pool was then subjected to mechanical and enzymatic disruption and further processing.

**Table 1.** Mock pool combinations comprising filarial nematodes and mosquitoes. Barcodes show sample number (i.e., 1-96) and quantity of mosquitoes in mock pool (i.e., 0M, 10M, 50M, or 100M) for a total of a 192 mock pools. L3 composition represents the corresponding filarial worms that have been spike into the pool with *Brugia malayi*, *Brugia pahangi*, and *Dirofilaria immitis*, respectively. The triplicate column differentiates repeated mock samples with a letter designation (i.e., A-C).

### Processing of mock pools

Mock pools were all subjected to the same DNA extraction protocol. Each sample was mechanically disrupted via the Omni International Bead Ruptor in a bead homogenizer tube with 2.8mm ceramic beads. After physical breakdown, further enzymatic disruption was done via proteinase K and tubes were placed thermomixer (56 °C; 350 rpm). Automated DNA extraction was accomplished in a Maxwell^®^ RSC Instrument using the Maxwell^®^ Tissue DNA Kit (Promega, Madison Wisconsin). All samples were then stored at -80 °C until further processing (Fig. 2).

### DNA preparation and sequencing

Amplicons of the mitochondrial cytochrome oxidase subunit I (*cox1*) gene of filarial nematodes amplified from the mock pools were sequenced by Oxford Nanopore Technologies (ONT) (Fig. 2). An initial PCR amplified a *cox1* fragment of ∼700bp with primers COINT (forward) 5’-TGATTGGTGGTTTTGGTAA-3’ and COINT (reverse) 5’-ATAAGTACGAGTATCAATATC-3’ using a modified protocol [35]. After initial PCR amplification, DNA amplicons were diluted to 200 fmol to prepare for Native Barcode ligation and to help remove PCR inhibitors. Each sample was ligated with a unique barcode (S1 Table) using a Native Barcoding Kit 96 V14 and carried out according to its specified protocol on the ONT website (SQK-NBD114.96). Pooled barcoded samples were ligated with a specific Native Adapter to aid DNA movement through nanopore for sequencing. After each ligation step, AMPure XP magnetic beads were used to clean up excess materials and a short fragment buffer was used with the final library being diluted to 35-50 fmol in 32 μL of elution buffer. The final library was added to the R10.4.1 flow cell (FLO-PRO114M) and loaded into a PromethION™ platform at the Molecular Genomics Core at the Texas A&M Institute of Genome Sciences and Society (TIGSS). Sequence data was made available via Texas A&M University’s Globus, a file transfer service designed for sharing data between data repositories for further downstream analyses.

### Bioinformatic analyses

In order to prepare sequences for bioinformatic analysis, raw signal data from Oxford Nanopore sequencing was analyzed using Dorado v0.8.2 with the super-accurate basecalling model v5.0.0 (dna_r10.4.1_e8.2_400bps_sup@v5.0.0). Reads were demultiplexed using dorado demux with the --barcode-both-ends option to retain only reads containing barcodes on both ends. Filtered reads were then clustered and consensus-called using *amplicon_sorter* (https://github.com/avierstr/amplicon_sorter) with the following parameters: -min 400 -max 800 -ar -np 48 -aln [36]. This configuration restricts clustering to reads between 400 and 800 bp to exclude off-target products, processes all reads within that size range, and outputs the multiple sequence alignment used to generate the consensus sequences. No more than 10,000 reads were processed per sample. All analyses were performed on the Grace computing cluster at Texas A&M University. The following command was used to download applicable sequences from NCBI’s GenBank “research -db nucleotide -query cox1[Gene] AND (Brugia OR Dirofilaria) AND mitochondrion[filter] | efetch -format fasta >all_filarial_cox1.fasta”.

Presence of filarial nematodes in each pooled sample was determined by sequenced and grouped reads. Negative control samples were used to help identify species-specific read thresholds (i.e., the highest species-specific read count in a negative control sample) (S2 Table) in order to determine the presence of filarial DNA within a sample [24, 37]. Blast analysis (NCBI) with default parameters identified the most closely matching filarial nematode species for each read group. Quantitative assessment of samples was determined by comparing the composition of filarial species in each mock pool with the corresponding number of sequence reads per L3 per species.

### Analysis of off-target reads

A small subset of samples (n=11) was processed as a quality control measure and to help compare single-end and double-end barcode data (Table 2). This included five samples re-sequenced and six samples re-extracted and subsequently sequenced.

**Table 2.** Reanalyzed mock pools. This table shows a sample subset (n=11) that were reanalyzed. Five mock pools were sequenced again (i.e., re-run) and six mock pools were re-extracted and subsequently sequenced (i.e., re-extraction). SB refer to single-end barcode results and DB refer to dual-ended barcode results. New Result (DB) refers to findings after mock pool was subjected to a re-run or re-extraction approach. Read counts for each pool can be found in the supporting information (S2 Table).

## 3. Results

### Relationship of B. malayi, B. pahangi, and D. immitis read numbers compared to the true mock species composition

A maximum of 10,000 reads were allowed to cluster for each pool of L3 filarial nematodes and mosquitoes. These were identified through BLAST analysis and reads were removed if they did not match sequences of one of the three filarial nematode species that were spiked into the mock samples based on the criteria >97% identity to whole read fragment. On mean average, 9,054 reads were retained in single-end barcode pools and 9,380 reads were retained in dual-end barcode pools that comprised at least one spiked filarial nematode. In addition, Bm averaged 5,103 reads for single-end barcode pools and 5,278 reads for dual-end barcode pools, Bp averaged and 3,365, and Di averaged 6,743 and 6,989, respectively (Fig. 3; S2 Table). However, these quantities varied widely depending on the species ratio of the spiked filarial nematodes. If, for example, the pool had a homogenous composition of filarial species (whether 1, 5 or 10 L3s of the same species), Bm counts had an average of 9,080 reads for single-barcode pools and 9492 reads for dual-end barcode pools, Bp averaged 8,479 and 8,792 reads, and Di averaged 9126 and 9466 reads, respectively (Fig. 3; S2 Table). Heterogenous pools (pools comprising three different filarial species in different ratios) did not have read counts that accurately compared to the true composition of worm numbers spiked into pools. However, 1:1:1 ratio (i.e., 1_1_1, 5_5_5, and 10_10_10) and 1:5:10 ratio (i.e., 1_5_10, 5_10_1, and 10_1_5) heterogenous pools showed similar averages despite this discrepancy. This included higher average read counts for Di pools and lower average read counts for Bp pools (Fig. 3; S2 Table).

**Figure 3.**
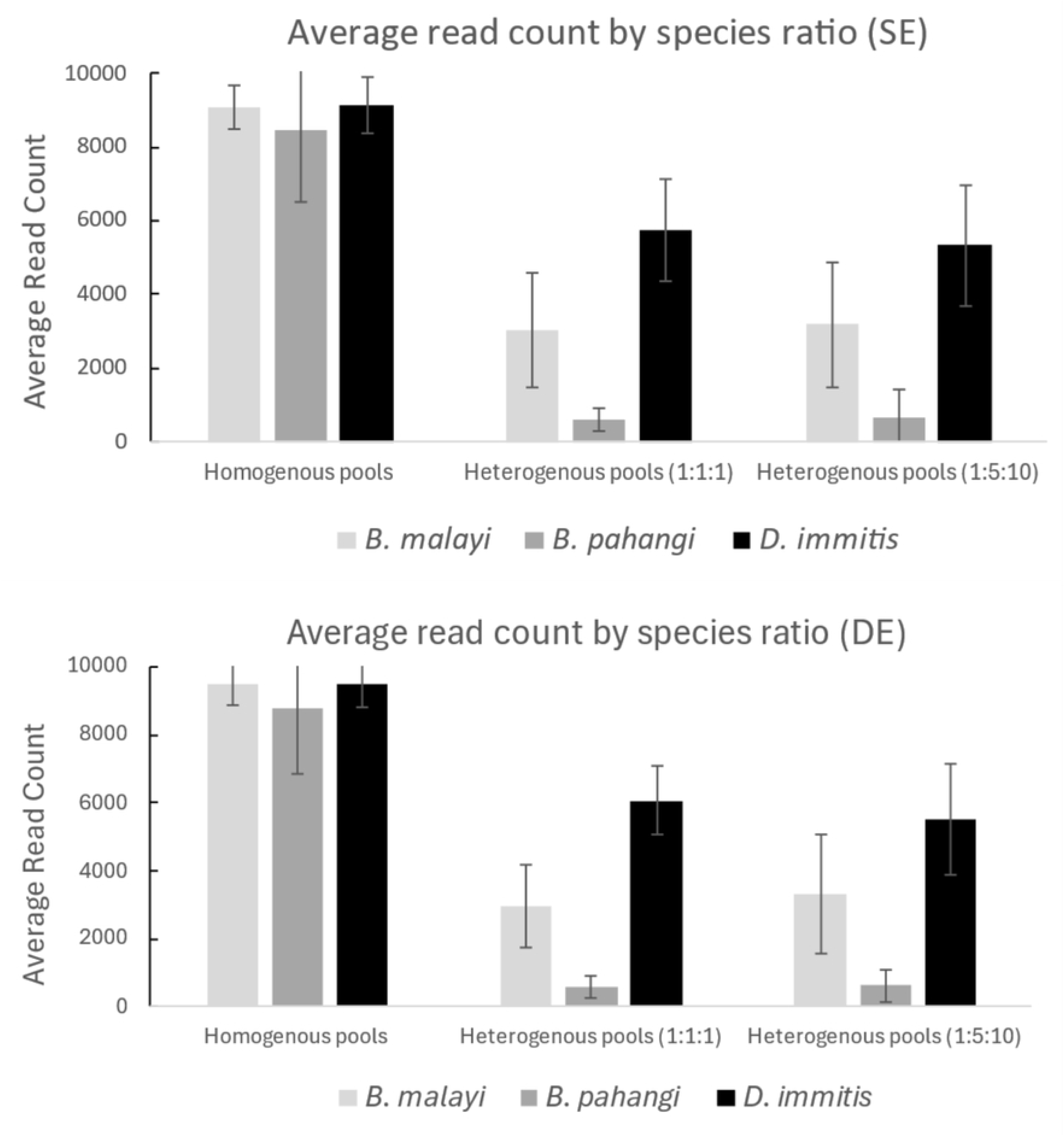
Average read counts based on different filarial nematode species ratios. The graph on top shows average read counts from single-end barcode reads (i.e., one barcode per fragment) and the graph on bottom shows double-end barcode reads (i.e., two barcodes per fragment). The bar graphs on the left represent homogenous pools, or pools with only on filarial species spiked in (in quantities of 1, 5, or 10). The center bar graphs show the average read counts in heterogenous pools of 1:1:1 ratio (i.e., 1_1_1, 5_5_5, and 10_10_10). The bar graphs on the right show the average read counts of heterogenous pools of 1:5:10 ratios (i.e., 1_5_10, 5_10_1, and 10_1_5). Filarial species are represented by the light gray (*B*. *malayi*), medium gray (*B*. *pahangi*), and dark gray (*D*. *immitis*).

### Relationship of Brugia malayi (Bm), Brugia pahangi (Bp), and Dirofilaria immitis (Di) ratios compared to the true mock species composition

Percentage species composition was calculated by taking the grouped reads per species and dividing them by the total number of reads from that sample/pool (S1; S2 Fig.). These calculations were greatly dependent on species identity of L3, the number of L3 spiked in, and the number of mosquito heads in the pool. If observing across all replicates and pools comprising all three filarial nematodes (using the dual-ended barcode approach), there was an average number of 33.11% (0.00-66.45%) Bm reads, 6.78% (0-31.59%) Bp reads, and 60.11% (26.61-95.47%) Di reads. When observing average species composition percentage of pools and replicates with all three species types (i.e., Bm, Bp, and Di), regardless of mosquito amounts, two patterns emerge depending on if they were 1:1:1 or 1:5:10 ratios.

Pools and replicates of 1:1:1 ratio were largely consistent despite the increase in the total amount of filarial nematodes spiked in. This included, respectively, average Bm reads of 27.76%, 31.74%, and 32.72%; Bp reads of 6.14%, 6.70%, and 6.07%; average Di reads of 66.10%, 61.57%, and 61.21%. Pools and replicates of 1:5:10 ratio were largely dependent the amount of each species that was spiked in. This included, respectively, average Bm reads of 15.41%, 45.75%, and 45.30%; Bp reads of 5.82%, 14.79%, and 1.18%; average Di reads of 78.78%, 39.46%, and 53.52%.

### Presence or absence of filarial species in mock communities using species-specific thresholds

Out of the 192 pooled samples that were initially sequenced and demultiplexed using a standard single-ended barcode approach, 93.75% of pools (n=180) accurately reflected the presence of the filarial nematode species composition (S2 Table). Out of these 12 samples that were inaccurate, 7 were false positive and 5 were false negative events. However, if we use species-specific threshold margins (Bm=178; Bp=85; Di=296 reads) detection of correct filarial composition decreased to 92.71% (n=178). These species-specific thresholds were created by using the maximum read count value for each filarial species in control mock pool samples (i.e., no spiked in filarial DNA). On the other hand, using a both-end barcode demultiplexing protocol allowed for 97.92% of pools (n=188) to correctly identify the presence of appropriate filarial DNA species (S2 Table). Out of these 4 samples that were inaccurate, two were false positive and two were false negative events. There were no changes using species-specific read thresholds which did not comprise any off-target reads in the control samples (Bm=0; Bp=0; Di=0 reads).

### Comparison of two demultiplexing pipelines

When using a single-ended demultiplexing pipeline, six of seven false positive pools were from samples comprising no filarial nematodes (e.g., 01-03_0M and 49-51_10M) and one was from a sample that only comprised Bp (65_10M) but those contaminated reads did not reach species-specific thresholds (Bm=34; Di=139). Ten false negative pools were spread among each of the four mosquito pool groups and included two in 0M (31_0M; 46_0M), three in 10M (94-96_10M), two in 50M (40-41_50M), and three in 100M (94-96_100M). Of those false negative pools, most were from missing Bp reads (n=7) and all were from pools that had only comprised a single L3 of a representative species spiked into a pool with multipleL3 of the two other species. A dual-ended demultiplexing pipeline comprised only one false positive from pools containing no filarial nematodes (50_10M) and the other, due to lower specific-species threshold, from Bp reads (65_10M). The two false negative pools were from missing Bp (95_10M) and Bm reads (41_50M).

A subset of samples was sequenced again (n=5) which comprised two correctly identified, two false positives, and one false negative when using a dual-ended barcode. After re-sequencing one true negative became a false positive and one false positive became a true negative (Table 2). In addition, a subset of samples was re-extracted and subsequently sequenced (n=6). Previously, five of these samples were correctly identified and only one was a false negative when using a dual-ended barcode. However, in the newly created mock pools, two correctly identified pools became false negative and the one false negative became true positive (Table 2).

## 4. Discussion

### Filariome diagnostic potential and molecular xenomonitoring

Accurate filarial species detection confirms the use of filariome as a tool to capture the full spectrum of diversity that is present within a tested sample, specifically, a tool for molecular xenomonitoring investigations. Demultiplexing with either a single-ended or dual-ended barcode showed 92.71% and 97.92% accuracy, respectively, for identifying correct filarial species presence. Furthermore, inaccuracies were most likely attributed to unnatural biases in the amplification targets. For example, many of the false negative pools were introduced when low DNA (i.e., one L3 of a certain species) was in the presence of high DNA amount (i.e., ten L3 of a certain species). These situations are unlikely to happen when molecular xenomonitoring because: i). multiple L3 worms would not likely be present in the arthropod vector head/mouth parts; and ii) the detection rate for filarial DNA in pools featuring 100 arthropod vector head/mouth parts is around 1% [4].

Having a highly accurate molecular approach to concomitantly detect multiple filarial nematodes species in a single pool/sample has critical implications to the fields of public health and veterinary medicine. As highlighted before, many filarial nematodes are NTDs or tropical diseases with devastating human health and economic impacts, including the agent of river blindness, the agents of lymphatic filariases, and the agent for loiasis. Molecular xenomonitoring is the primary diagnostic tool for active prevention, elimination, and surveillance efforts. Therefore, preventing data loss through pan-filarial high throughput sequencing (i.e., filariome), could help mitigate and thwart further human suffering.

Moreover, these principles can extend to filarial nematodes causing harm in both domestic and wild animals due to increased animal movement and climate change [16, 19, 34, 36]. For example, when collecting black flies to monitor the presence of *O*. *volvulus* in sub-Saharan Africa, one could also acquire data about the sympatric transmission of *O*. *ochengi*. While the life cycles of filarial nematodes of public health and veterinary relevance are generally well-understood, much remains to be elucidated on the biology of filarial nematodes infecting wildlife or that are recently emerging [17, 20, 38, 39]. As the chance for spillover infections increase due to anthropogenic climate change and habitat degradation [40–42], it becomes pertinent to surveillance for understudied filarial nematode biodiversity, their biogeographic distribution, and host-vector-parasite assemblages. It is also increasingly likely that uncharacterized cryptic filarial species and unknown intraspecies variability, could alter the dynamics of assemblages and host health [20, 21]. This includes the potential for some filarial populations to become resistant to ivermectin and threaten MDA capabilities [43]

Apart from the data lost during targeted molecular xenomonitoring, remote and laborious working conditions with high cost remain some of the largest obstacles to this type of surveillance. However, using nanopore sequencing (ONT) circumvents these challenges by employing a higher throughput, portability, and providing a cheaper alternative to other molecular technologies such as the Illumina platform [44–46]. In addition to this versatility, the filariome approach offers clinical capabilities at higher taxonomic resolution (i.e., longer reads). Nanopore sequencing using the MinION™ device has been previously used for screening blood of domestic dogs for filarial nematodes [32]. Indeed, one could extrapolate the use of Filariome to test both humans and animals for co-infections by two or more filarial nematode species using their blood or other biological samples, such as skin biopsies. Having a comprehensive, single approach, regardless of geographic or social context, would allow global coordination of filarial worm diagnosis and elimination efforts.

### Quantitative measurements and limitations

Biases in species specific read counts were persistent throughout the experiment, in particular, the overrepresentation of Di reads and the underrepresentation of Bm and Bp reads. For example, when looking at pools with a 1:1:1 ratio of the three spiked filarial species, Bm reads were on average half and Bp reads were on average a tenth of Di reads. While these biases seem to exist in other nematode deep amplicon sequencing analyses [24, 32], it makes interpretation of quantitative data difficult. Correction factors can diminish these effects, but this study did not or could not show they were helpful to data interpretation. A larger replicate sample size for pools (n>3) would be ideal in this experiment to justify a consistent calculation. Regardless, correction factors were calculated out of all pools with 1:1:1 and 1:5:10 ratio comprising no mosquitoes. Correction factors were largely variable and did not prove useful for determining expected read counts (S3 Table) At this moment, it remains unclear why there is an over-and underrepresentation in read counts for certain species, but there are various possible explanations. For one, L3 sizes can differ among filarial species, and thus provide more cells, and thus DNA, to be amplified. However, morphological data (e.g., length and width) show that these species are of relatively the similar dimensions [47–51], so this idea is less likely to be accurate. In addition, differences in morphology, such as the presence of a sheath in species of *Brugia* but absent in Di, could limit the effectiveness of DNA lysis. Future work would be prudent to employ more than one DNA extraction technique simultaneously to optimize the protocol. Another reasonable explanation for the species-specific read differences is from copy number variations. While nuclear DNA (nDNA) only possess two copies of a gene per cell, mitochondrial DNA (mtDNA) can have high numbers of copies and vary greatly between filarial species [52, 53]. Lastly, interspecies variability may impact the amplication efficiency or the primer binding site. If that were the case, it would explain why some filarial worms (e.g., Bp) appear to be at a competitive disadvantage to be amplified compared to others (e.g., Bm, Di). It’s also important to note that these were identically bred lab strains of each species. Intraspecies variability is likely to play some role in primer binding efficiency.

### Downstream analysis and contamination

The use of single-end or double-end barcodes had a sizable impact on the downstream analysis. We saw a difference of 5.21% accuracy between these methods, or ten sample pools that could be correctly or incorrectly identified. Thus, it is strongly recommended for future studies to use a double-ended barcode during bioinformatic analysis to reduce barcode hopping and increase both sensitivity and specificity. We also ran a subset of 11 samples (Table 2) to ensure low read counts inducing false positives/negatives are a common finding requiring filtering and not the result of contamination during DNA processing. Interestingly, five samples became the opposite of their former result. These findings support that these off-target reads are not the product of human or laboratory cross-contamination, but rather a common by-product of nanopore sequencing. While these off-target reads are not an uncommon finding [24, 32], this study highlights the importance of the type of downstream analysis used and the creation of read thresholds. In an actual surveillance event, false negatives would rarely be identified because there would be no further downstream analysis. However, these false negatives scenarios seem to only occur in pools that have heterogenous species composition with a mixed ratio (1:5:10). These extreme DNA concentration differences, which lead to false negative events, would be unlikely to occur when molecular xenomonitoring, but it does reveal the maximum limits of this tool. It should be noted that since certain filarial species have under-and overrepresentation of read counts, we utilized species-specific read thresholds even though it does not appear to be a common practice in the literature.

## Acknowledgments

The following reagents were provided by the NIH/NIAID Filariasis Research Reagent Resource Center for distribution through BEI Resources, NIAID, NIH: Uninfected *Aedes aegypti*, Strain Black Eye Liverpool (Frozen), NR-48920; Stage L3 *Brugia malayi* Infective Larvae (Frozen), NR-48891; Stage L3 *Brugia pahangi* Infective Larvae (Frozen), NR-48900; Stage L3 *Dirofilaria immitis*, Strain Missouri, Infective Larvae (Frozen), NR-48910.

## Availability of data and material

All data generated or analyzed during this study are included in this published article.

## Authors’ contributions

MRK and GGV conceptualized the study. MRK and GGV drafted the manuscript. JSG and GGV: supervision. MRK and GGV acquired samples. MRK and CB performed molecular and bioinformatic analysis. All authors read and approved the final manuscript.

## Ethics approval

Not applicable

## Supporting Information

**S1 Fig. Percentage composition of reads for each mock pool using single-end barcodes.**

Bar graphs depicting the percentage of reads from each filarial species (Bm=gray; Bp=blue; Di=red) broken down by individual mock pools (x-axis). Each panel shows mock pools comprising different amounts of mosquitoes (i.e., 0-top left, 10-top right, 50-bottom left, 100-bottom right). This data was analyzed using single-end barcodes.

**S2 Fig. Percentage composition of reads for each mock pool using double-end barcodes.**

Bar graphs depicting the percentage of reads from each filarial species (Bm=gray; Bp=blue; Di=red) broken down by individual mock pools (x-axis). Each panel shows mock pools comprising different amounts of mosquitoes (i.e., 0-top left, 10-top right, 50-bottom left, 100-bottom right). This data was analyzed using double-end barcodes.

**S1 Table. Unique Barcodes used for deep amplicon sequencing**

Each sample was ligated with a unique barcode from a Native Barcoding Kit 96 V14 kit. The protocol used for barcode ligation can be found at the Oxford Nanopore website (SQK-NBD114.96).

**S2 Table. Filarial species read counts in each mock pool**

Each tab comprises mock pools from 0, 10, 50, or 100 mosquitoes (i.e., 0M. 10M, 50M, or 100M) and analyzed with single-end (SE) or double-end (DE) barcodes. Read counts are broken down by filarial species and their mock pool label. Mock pools that have missing or off-target reads are highlighted (see footnotes). In addition, the average read count for each filarial species across replicates is noted.

**S3 Table. Determining the usage of correction factors**

To calculate for correction factors, we used the mathematical formula from previous nemabiome work [24]. This involves dividing the true proportion of the filarial species in the pool (based on counted L3s) by the mean proportion of nanopore sequences generated from the three replicates (correction factor = %Actual / %Observed). This was done in a variety of scenarios from pools with 0M. Each scenario would largely produce unique correction factors rather than a single consistent value across pools.

## References

1. Anderson RC. Nematode parasites of vertebrates: Their Development and Transmission: CABI, Wallingford, Oxon, UK; 2000.

2. Gyasi ME, Okonkwo ON, Tripathy K. Onchocerciasis. StatPearls [Internet]: StatPearls Publishing; 2023.

3. World Health Organization. Lymphatic filariasis. https://www.who.int/news-room/fact-sheets/detail/lymphatic-filariasis 2024 [Accessed on November 2024]. Available from: https://www.who.int/news-room/fact-sheets/detail/lymphatic-filariasis.

4. Pilotte N, Unnasch TR, Williams SA. The current status of molecular xenomonitoring for lymphatic filariasis and onchocerciasis. Trends Parasitol. 2017;33(10):788–98.

5. World Health Organization. Guidelines for stopping mass drug administration and verifying elimination of human onchocerciasis: World Health Organization; 2016 [cited 2024 Accessed August 7 ].

6. Rao RU, Samarasekera SD, Nagodavithana KC, Punchihewa MW, Dassanayaka TD, PK D G, et al. Programmatic use of molecular xenomonitoring at the level of evaluation units to assess persistence of lymphatic filariasis in Sri Lanka. PLoS Negl Trop Dis. 2016;10(5):e0004722.

7. Lau CL, Won KY, Lammie PJ, Graves PM. Lymphatic filariasis elimination in American Samoa: evaluation of molecular xenomonitoring as a surveillance tool in the endgame. PLoS Negl Trop Dis. 2016;10(11):e0005108.

8. Lakwo T, Oguttu D, Ukety T, Post R, Bakajika D. Onchocerciasis elimination: progress and challenges. Res Rep Trop Med. 2020:81–95.

9. World Health Organization. Certification of elimination of human onchocerciasis: criteria and procedures 2001.

10. World Health Organization. Monitoring and epidemiological assessment of mass drug administration in the global programme to eliminate lymphatic filariasis: a manual for national elimination programmes 2011.

11. Small ST, Labbé F, Coulibaly YI, Nutman TB, King CL, Serre D, et al. Human migration and the spread of the nematode parasite *Wuchereria bancrofti*. Mol Biol Evol. 2019;36(9):1931–41.

12. Gardon J, Gardon-Wendel N, Kamgno J, Chippaux J-P, Boussinesq M. Serious reactions after mass treatment of onchocerciasis with ivermectin in an area endemic for *Loa loa* infection. The Lancet. 1997;350(9070):18–22.

13. Boussinesq M, Gardon J, Gardon-Wendel N, Chippaux J-P. Clinical picture, epidemiology and outcome of *Loa*-associated serious adverse events related to mass ivermectin treatment of onchocerciasis in Cameroon. Filaria J. 2003;2(Suppl 1):S4.

14. Anderson RC. Filarioid Nematodes. Parasitic Diseases of Wild Mammals. 2001:342–56.

15. Nelson GS. The pathology of filarial infections. Helminthol Abstr. 1966; 35:311–336

16. Verocai GG, Gomez JL, Hakimi H, Kulpa MR, Luksovsky JL, Thompson DP, et al. Validation of a species-specific probe-based qPCR for detection of *Setaria yehi* (Filarioidea: Onchocercidae) in Alaskan moose (*Alces alces gigas*). Int J Parasitol Parasites Wildl. 2024;25:100990.

17. Gruntmeir J, Kelly M, Ramos RAN, Verocai GG. Cutaneous filarioid nematodes of dogs in the United States: Are they emerging, neglected, or underdiagnosed parasites? Frontiers VetSci. 2023;10:1128611.

18. Orihel TC, Eberhard ML. Zoonotic filariasis. Clin Microbiol Rev. 1998;11(2):366–81.

19. Lefoulon E, Giannelli A, Makepeace BL, Mutafchiev Y, Townson S, Uni S, et al. Whence river blindness? The domestication of mammals and host-parasite co-evolution in the nematode genus *Onchocerca*. Int J Parasitol. 2017;47(8):457–70. Epub 2017/03/28. doi: 10.1016/j.ijpara.2016.12.009. PubMed PMID: 28344097.

20. Kulpa MR, Lefoulon E, Beckmen KB, Allen SE, Malmberg J, Crouse JA, et al. A footworm in the door: revising *Onchocerca* phylogeny with previously unknown cryptic species in wild North American ungulates. Int J Parasitol. 2025;55(1):59–68.

21. Cháves-González LE, Morales-Calvo F, Mora J, Solano-Barquero A, Verocai GG, Rojas A. What lies behind the curtain: Cryptic diversity in helminth parasites of human and veterinary importance. Curr Res Parasitol Vector Borne Dis. 2022;2:100094.

22. Colella V, Young ND, Manzanell R, Atapattu U, Sumanam SB, Huggins LG, et al. *Dirofilaria asiatica* sp. nov. (Spirurida: Onchocercidae)–Defined using a combined morphological-molecular approach. Int J Parasitol. 2025; 55:461–474.

23. Benedict BM, Barboza PS, Crouse JA, Groch KR, Kulpa MR, Thompson DP, et al. Sores of boreal moose reveal a previously unknown genetic lineage of parasitic nematode within the genus *Onchocerca*. PLoS One. 2023;18(1):e0278886.

24. Avramenko RW, Redman EM, Lewis R, Yazwinski TA, Wasmuth JD, Gilleard JS. Exploring the gastrointestinal “nemabiome”: deep amplicon sequencing to quantify the species composition of parasitic nematode communities. PLoS One. 2015;10(12):e0143559.

25. Beaumelle C, Redman E, Verheyden H, Jacquiet P, Bégoc N, Veyssière F, et al. Generalist nematodes dominate the nemabiome of roe deer in sympatry with sheep at a regional level. Int J Parasitol. 2022;52(12):751–61.

26. Halvarsson P, Baltrušis P, Kjellander P, Höglund J. Parasitic strongyle nemabiome communities in wild ruminants in Sweden. Parasit Vectors. 2022;15(1):341.

27. Queiroz C, Levy M, Avramenko R, Redman E, Kearns K, Swain L, et al. The use of ITS-2 rDNA nemabiome metabarcoding to enhance anthelmintic resistance diagnosis and surveillance of ovine gastrointestinal nematodes. Int J Parasitol Drugs Drug Resist. 2020;14:105–17.

28. Courtot É, Boisseau M, Dhorne-Pollet S, Serreau D, Gesbert A, Reigner F, et al. Comparison of two molecular barcodes for the study of equine strongylid communities with amplicon sequencing. PeerJ. 2023; 11:e15124.

29. Chelladurai JRJ, Johnson WL, Quintana TA, Nilaweera GW, Wolfe H, Wehus-Tow B, et al. Nemabiome metabarcoding to assess the diversity of trichostrongyle nematodes in plains bison from the mid-western USA. Preprint. 10.21203/rs.3.rs-4632804/v1.

30. Avramenko RW, Redman EM, Lewis R, Bichuette MA, Palmeira BM, Yazwinski TA, et al. The use of nemabiome metabarcoding to explore gastro-intestinal nematode species diversity and anthelmintic treatment effectiveness in beef calves. Int J Parasitol. 2017;47(13):893–902.

31. Kipp KR, Redman EM, Luksovsky JL, Claussen D, Gilleard JS, Verocai GG. High frequency of benzimidazole resistance polymorphisms and age-class differences in trichostrongyle nematodes of ranched bison from the south-central United States. Int J Parasitol Drugs Drug Resist. 2025;28:100594.

32. Huggins LG, Atapattu U, Young ND, Traub RJ, Colella V. Development and validation of a long-read metabarcoding platform for the detection of filarial worm pathogens of animals and humans. BMC Microbiol. 2024;24(1):28.

33. World Health Organization. Defining the roles of vector control and xenomonitoring in the Global Programme to Eliminate Lymphatic Filariasis: report of the informal consultation WHO/HQ, Geneva, 29-31 January 2002 2002.

34. Michalski ML, Griffiths KG, Williams SA, Kaplan RM, Moorhead AR. The NIH-NIAID filariasis research reagent resource center. PLoS Negl Trop Dis. 2011;5(11):e1261.

35. Casiraghi M, Anderson T, Bandi C, Bazzocchi C, Genchi C. A phylogenetic analysis of filarial nematodes: comparison with the phylogeny of *Wolbachia* endosymbionts. Parasitology. 2001;122(1):93–103.

36. Vierstraete AR, Braeckman BP. Amplicon_sorter: A tool for reference-free amplicon sorting based on sequence similarity and for building consensus sequences. Ecol Evol. 2022;12(3):e8603.

37. Early AM, Daniels RF, Farrell TM, Grimsby J, Volkman SK, Wirth DF, et al. Detection of low-density *Plasmodium falciparum* infections using amplicon deep sequencing. Malaria J. 2019;18:1–13.

38. Kulpa M, Goldsmith D, Verocai GG. An unusual case of *Brugia* sp. infection in a dog from Alberta, Canada. Vet. Parasitol. Reg. Stud. .Rep.. 2022;37:100811.

39. Grácio AJS, Richter J, Komnenou AT, Grácio MA. Onchocerciasis caused by *Onchocerca lupi*: an emerging zoonotic infection. Systematic review. Parasitol Res. 2015;114(7):2401–13.

40. Laaksonen S, Pusenius J, Kumpula J, Venäläinen A, Kortet R, Oksanen A, et al. Climate change promotes the emergence of serious disease outbreaks of filarioid nematodes. EcoHealth. 2010;7(1):7–13.

41. Gordon CA, McManus DP, Jones MK, Gray DJ, Gobert GN. The increase of exotic zoonotic helminth infections: The impact of urbanization, climate change and globalization. Adv Parasitol. 2016;91:311–97.

42. Hoberg EP, Brooks DR. Evolution in action: climate change, biodiversity dynamics and emerging infectious disease. Philos Trans Royal Soc B. 2015;370(1665):20130553.

43. Harshita A, Nonika R. Emerging antihelminthic drug resistance: Implications for mass drug administration program. One Health Bull. 2024;4(4):157–63.

44. Quick J, Loman NJ, Duraffour S, Simpson JT, Severi E, Cowley L, et al. Real-time, portable genome sequencing for Ebola surveillance. Nature. 2016;530(7589):228–32.

45. Chen P, Sun Z, Wang J, Liu X, Bai Y, Chen J, et al. Portable nanopore-sequencing technology: Trends in development and applications. Front Microbiol. 2023;14:1043967.

46. Menegon M, Cantaloni C, Rodriguez-Prieto A, Centomo C, Abdelfattah A, Rossato M, et al. On site DNA barcoding by nanopore sequencing. PLoS One. 2017;12(10):e0184741.

47. Mutafchiev Y, Bain O, Williams Z, McCall JW, Michalski ML. Intraperitoneal development of the filarial nematode *Brugia malayi* in the Mongolian jird (*Meriones unguiculatus*). Parasitol Res. 2014;113(5):1827–35.

48. Schacher JF. Developmental stages of *Brugia pahangi* in the final host. J Parasitol. 1962; 48:693–706.

49. Orihel TC. Morphology of the larval stages of *Dirofilaria immitis* in the dog. J Parasitol. 1961;47(2):251–62.

50. Hess JA, Eberhard ML, Segura-Lepe M, Grundner-Culemann K, Kracher B, Shryock J, et al. A rodent model for *Dirofilaria immitis*, canine heartworm: parasite growth, development, and drug sensitivity in NSG mice. Sci Rep. 2023;13(1):976.

51. Ash LR, Riley JM. Development of subperiodic *Brugia malayi* in the jird, *Meriones unguiculatus*, with notes on infections in other rodents. J Parasitol. 1970:969–73.

52. Laidoudi Y, Davoust B, Varloud M, Niang EHA, Fenollar F, Mediannikov O. Development of a multiplex qPCR-based approach for the diagnosis of *Dirofilaria immitis*, D. repens and Acanthocheilonema reconditum. Parasit Vectors. 2020;13(1):1–15.

53. Papaiakovou M, Waeschenbach A, Ajibola O, Ajjampur SS, Anderson RM, Bailey R, et al. Global diversity of soil-transmitted helminths reveals population-biased genetic variation that impacts diagnostic targets. Naturee Comm. 2025:16:6374.

